# Structural insights into tick-borne encephalitis virus neutralization and animal protection by a therapeutic antibody

**DOI:** 10.1101/2021.07.28.453943

**Authors:** Ivan K. Baykov, Grzegorz Chojnowski, Petr Pachl, Andrey L. Matveev, Nina A. Moor, Lyudmila A. Emelianova, Pavlina M. Rezacova, Victor S. Lamzin, Nina V. Tikunova

## Abstract

Tick-borne encephalitis virus (TBEV) causes about 5-6 thousand cases annually, while there is still no effective treatment for this virus. To fill this gap, a high-affinity chimeric anti-TBEV antibody ch14D5 has previously been constructed, and high protective activity in a murine TBEV model has been shown for this antibody. However, the mechanism of action of this antibody and the recognized epitope have not been known yet. In this study, it is shown by X-ray crystallography that this antibody recognizes a unique epitope on the lateral ridge of the D3 domain of glycoprotein E, which is readily accessible for binding. The orientation of this antibody relative to the virion surface makes bivalent binding possible, which facilitates the cross-linking of glycoprotein E molecules and thus blocking of surface rearrangements required for infection. Since the antibody tightly binds to this protein even at pH ∼ 5.0, it locks the virion in an acidic environment inside the late endosomes or phagosomes and, therefore, effectively blocks the fusion of the viral and endosomal/phagosomal membranes. We believe that this is why the ch14D5 antibody does not induce an antibody-dependent enhancement of infection *in vivo*, which is critical in the development of antibody-based therapeutic agents. In addition, the structure of the antibody-glycoprotein E interface can be used for the rational design of this antibody for enhancing its properties.

## Introduction

The tick-borne encephalitis virus (TBEV) belongs to tick-transmitted flaviviruses, and causes 5−6 thousand cases of the disease annually, frequently accompanied by meningitis, meningoencephalitis, or encephalomyelitis (Ruzek D et al., 2019; ECDC 2021 report). Approximately two thousand cases are registered in Russia (Nikitin AYa et al., 2020). There are three main subtypes of TBEV – the Far-Eastern, Siberian, and European, and the mortality caused by the virus subtypes ranges from 1−2% for the European subtype to 5−20% for the Far-Eastern subtype (Mansfield KL et al., 2009). Currently, there are no approved therapeutic agents to treat TBE (Elsterova J et al., 2017). Despite the fact that effective vaccines against TBEV have been developed for adults and children, the vaccination rate in European countries and Russia remains insufficient. It is estimated that approximately 25% of the population is vaccinated against TBEV in Europe, whereas less than 10%, in Russia (Erber W et al., 2018; Nikitin AYa et al., 2020).

Recombinant antibodies are one of the promising types of medicines against flaviviruses, and TBEV in particular. However, the therapeutic efficacy of a particular antibody depends on many factors, such as the affinity of the antibody, the location of the recognized epitope on the surface of the viral protein and its spatial availability, the geometric features of antibody binding, and the involvement of various mechanisms of the immune system (Pierson TC et al., 2008; Dowd KA et al., 2011; Dai L et al., 2016; Sun H et al., 2018). These factors also affect whether a particular antibody is prone to cause undesirable antibody-mediated enhancement (AME) of infection or not (Pierson TC et al., 2007; Pierson TC et al., 2008; Dowd KA et al., 2011). Several neutralizing or protective monoclonal antibodies have previously been described against TBEV (Guirakhoo F et al., 1989; Mandl −CW et al., 1989; Holzmann H et al., 1995; Tsekhanovskaya NA et al., 1993; Levanov LN et al., 2010; Kiermayr S. et al., 2009; Matveev AL et al., 2019; Agudelo M et al., 2021). However, the 3D structures of the antigen-antibody complex have only been published for three of them: 19/1786, Mab 4.2 and T025 (Niedrig M et al. 1994; Fuzik T et al., 2018; Jiang W et al., 1994; Yang X et al., 2019; Agudelo M et al., 2021). These antibodies are virus-neutralizing and bind adjacent epitopes on the lateral ridge of the D3 domain of glycoprotein E. This region is believed to interact with cell receptors and mediate virus binding, supported by the fact that the most potent virus-neutralizing mouse antibodies against various flaviviruses bind this ridge (Mandl CW, 2005; Kellman EM et al., 2018).

Another promising antibody against TBEV, named ch14D5, has previously been constructed and studied (Baykov IK et al., 2014). This antibody binds to the D3 domain of TBEV glycoprotein E (Sofjin-Ru strain) with a nanomolar affinity and also provides protection of mice infected with high lethal doses of the TBEV strain Absettarov. Moreover, this antibody caused no detectable AME in mice (Baykov IK et al., 2014; Baykov IK et al., 2018), which is very important for the development of antibody-based therapeutic agents. Given such prominent antiviral properties, this antibody is considered as a potential therapeutic agent for the prevention and treatment of TBE. However, the detailed structure of the recognized epitope, as well as the binding geometry and molecular mechanism of this antibody still remain unclear.

In this study, we report the X-ray structure of the complex formed by a Fab fragment of the ch14D5 antibody and the D3 domain of TBEV glycoprotein E, Sofjin-Ru strain (Far-Eastern subtype), determined at 2.3 Å resolution. Analysis of the interface between the proteins revealed key amino acid residues comprising the epitope and involved in complex formation. Binding orientation significantly differs for the ch14D5 antibody and other anti-TBEV antibodies with a known structure, while it still allows for bivalent antibody binding. It was shown that the antibody-antigen complex is stable at pH 4.8, which is critical for the ability of antibody to trap virion during the acidification of the inner space of endosomes. Based on the data, an antiviral mechanism for the ch14D5 antibody was proposed, which explains the high antiviral properties of this antibody and the absence of antibody-dependent enhancement of infection *in vivo*.

## Results

### X-ray crystal structure of the ch14D5 antibody-D3 domain complex, and the analysis of the recognized epitope

The structure of the ch14D5 antibody Fab fragment complex co-crystallized with the D3 domain of TBEV glycoprotein E, Sofjin-Ru strain, was determined by X-ray diffraction. As far as we know, this is the first published structure containing the D3 domain from the Far-Eastern TBEV subtype containing an alanine residue at position 331. The structure was determined at 2.3 Å resolution sufficient to analyze the fine structure of the antibody-antigen interface (pdb id NNNN, *currently in deposition process*) (Figure 1A, Table S1). The contacting amino acid residues of the D3 domain form two regions: residues D308 – K315 (an A_x_A loop of the D3 domain and a part of an A strand) and residues A331 – K336 (a BC_x_ loop and a part of a B strand) (Table S2). The numbering scheme and loop names were chosen according to the flavivirus glycoprotein E numbering (Rey FA et al., 1995). Amino acid residues N367 and E387 located in a D_x_E and a FG loops can also participate in complex formation. The interface is primarily stabilized by electrostatic interactions and direct or water-bridged hydrogen bonds, as well as by hydrophobic interactions to some extent. The angle between the longitudinal axis of the Fab fragment and virion surface is about 40° (Figure 1B), and the total buried surface area of the interface is approximately 670 Å^2^. All contacting residues of the D3 domain are localized within its lateral ridge. Since this part of the D3 domain is believed to be involved in interaction with cell receptors (Pierson TC et al., 2008; Sun H et al., 2018; Laureti M et al., 2018), we consider that the blocking of virion binding to cell receptors is one of the major mechanisms of the antiviral activity of the ch14D5 antibody.

**Figure 1.**
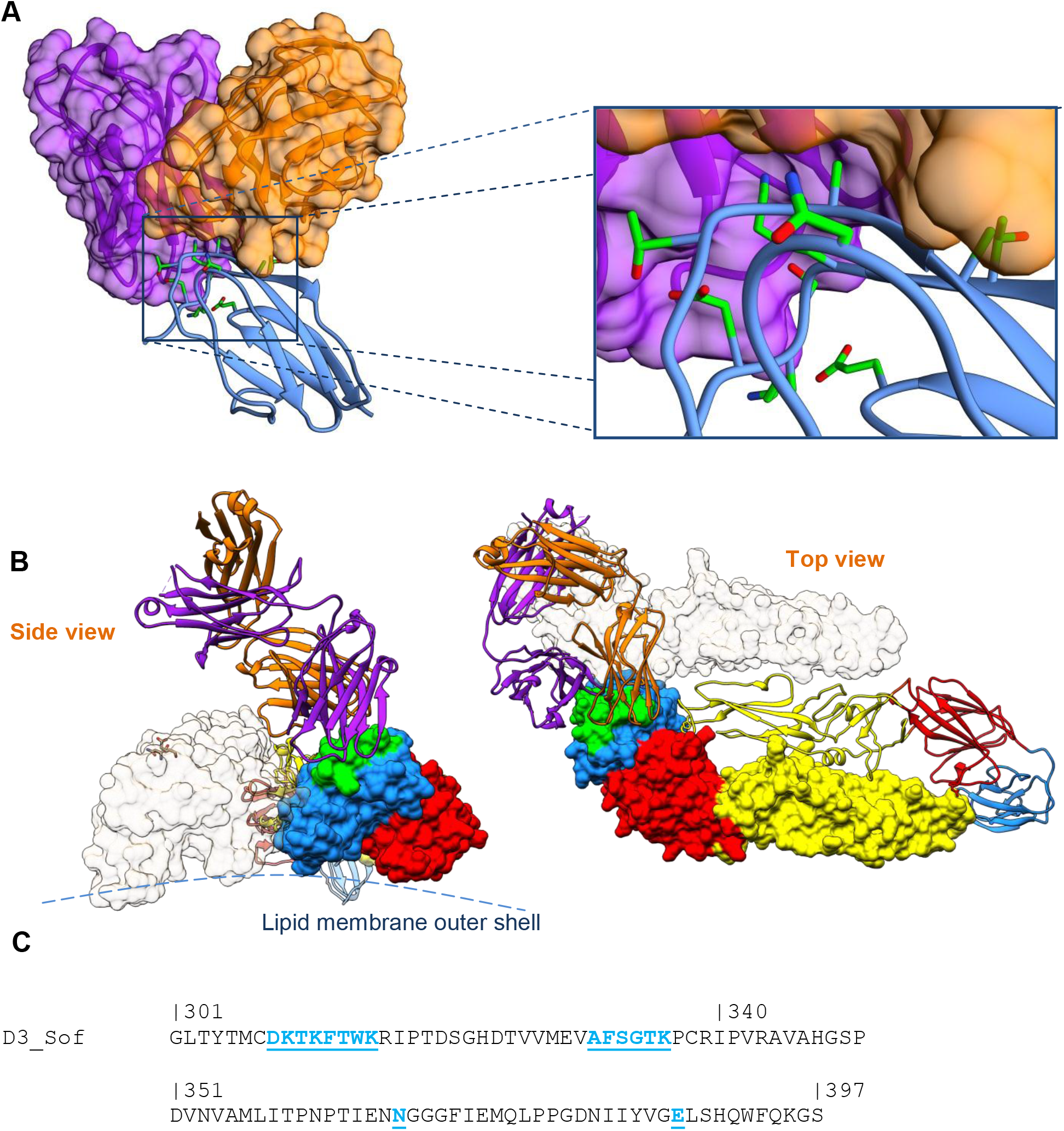
Three-dimensional structure of the ch14D5 antibody Fab fragment bound to the D3 domain of glycoprotein E. (A) Overall complex structure (only the Fv fragment of antibody is shown) and detailed interface (within inset). Heavy and light chains are shown in purple and orange, and the D3 domain is shown in blue. Residues forming contacts are in neon-green. (B) Structure of the complex within mature virion surface context. Domain D1, D2 and D3 are shown in red, yellow and blue, respectively. Residues forming the lateral ridge are shown in green. (C) Amino acid sequence of domain D3 with epitope residues is shown in blue and underlined.

### The effect of the D3 domain amino acid variations on the ch14D5 antibody binding

The D3 domain of TBEV glycoprotein E is highly conserved, however there are some subtype-specific amino acid variations at positions 313, 317 and 331 (Figure S1; Ecker M et al., 1999). We analyzed the effect of these variations on the ch14D5 antibody binding. For this analysis, we used recombinant D3 proteins based on cDNA fragments of Sofjin-Ru strain (Far-Eastern subtype of TBEV), Zausaev-like and Vasilchenko-like strains (Siberian subtype of TBEV) and Absettarov strain of TBEV (European subtype) as well as strains of Baltic and Bosnia lineages of Siberian subtype of TBEV and one strain of Omsk hemorrhagic fever virus (OHFV) (Figure S2A). These proteins differ at positions 313 and 331, which are within the epitope, as well as at position 317, which is outside the epitope according to the X-ray structure (Figure S2B). Using surface plasmon resonance (SPR) technique, it was shown that the ch14D5 antibody exhibited the highest affinity (K_D_ = 3.0 ± 1.4 nM) for the D3_Sof protein, while the affinities for other D3 proteins were substantially weaker (Figures 2 and S3, Table S3). There still was a minor possibility that the differences in synonymous codons between the same positions of different D3 genes may affect the rate of protein synthesis and, hence, the folding of the D3 proteins leading to differences in affinity. To rule out this possibility, we obtained a mutant protein called D3_SofA331T, which was coded by D3_Sof gene containing “GCG” to “ACC” mutation encoding Ala331Thr substitution. As expected, the affinity of the ch14D5 antibody for the mutant variant was noticeably weaker than for the D3_Sof protein, which confirmed the importance of the residue at position 331 of the glycoprotein E D3 domain for the ch14D5 antibody binding.

**Figure 2.**
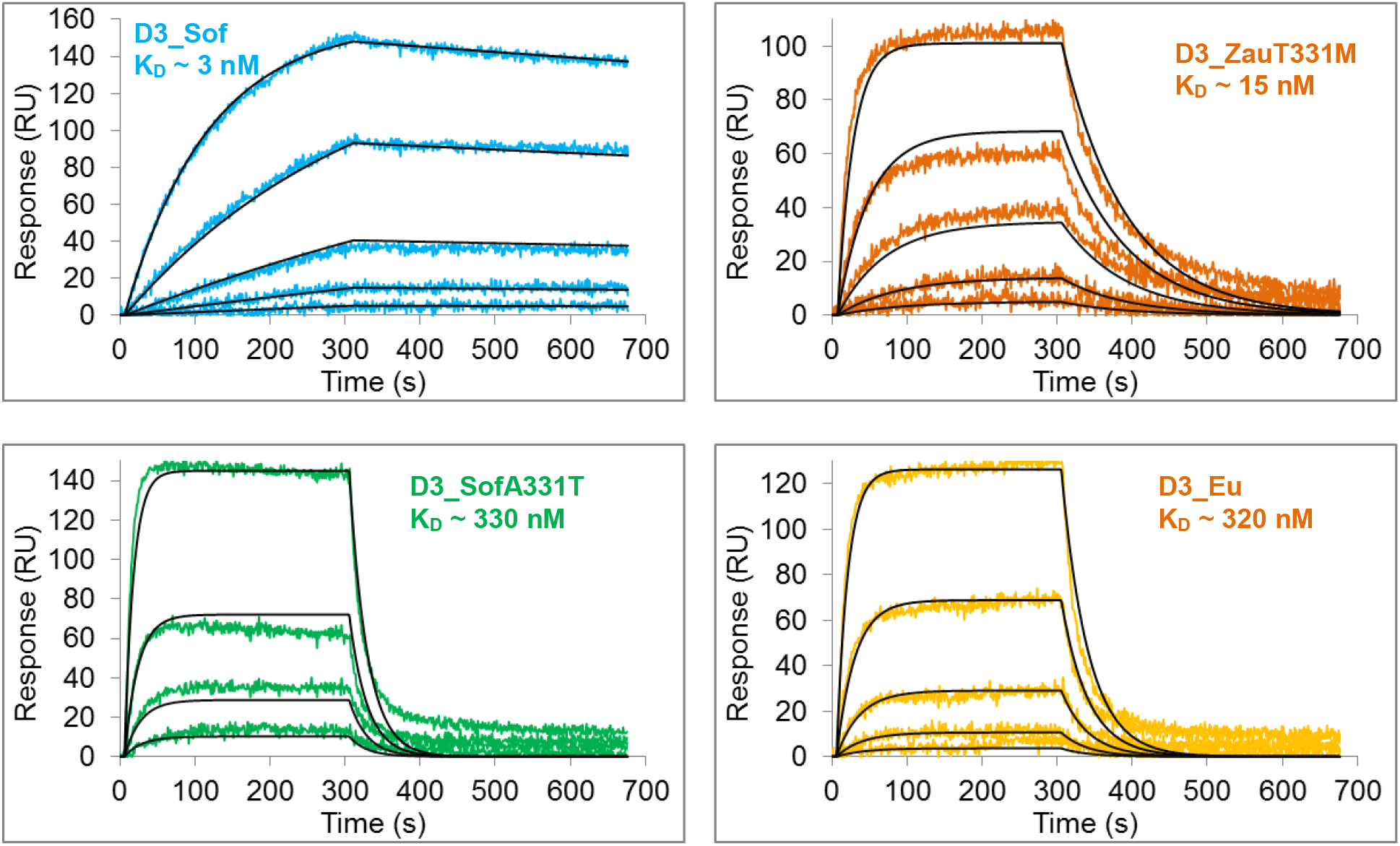
Differences in binding of the ch14D5 antibody to recombinant D3 proteins. Binding of the ch14D5 antibody to recombinant proteins D3_Sof, D3_Eu, D3_ZauT331M and D3_SofA331T examined by SPR. The D3 proteins were immobilized onto biosensor chip surface, the ch14D5 antibody was used as three-fold dilutions starting from 81 nM concentration (in the case of D3_Sof and D3_ZauT331M proteins) or from 405 nM (in the case of D3_Eu and D3_SofA331T proteins). Experimental curves are shown by colored lines, approximations are shown in black. The analysis was carried out using a ProteOn XPR36 biosensor. See also Figure S3 and Table S3.

### Accessibility of the ch14D5 epitope on the surface of the TBEV virion for monovalent and bivalent binding

The surface of the mature TBEV virion contains 180 glycoprotein E molecules (Rey FA et al., 1995). Due to the icosahedral symmetry of the virion, there are three different types of glycoprotein E molecules which differ in the accessibility of certain epitopes (Kaufmann B et al., 2006). These types are commonly marked as A, B and C regarding their proximity to 5-, 2- and 3-fold symmetry axes, respectively (Figure 3A). Since the accessibility of the recognized epitope and the number of the accessible epitope copies significantly affects the virus-neutralizing potency of a certain antibody (Kaufmann B et al., 2006), we analyzed the accessibility of the epitope recognized by the ch14D5 antibody on the surface of virion. A model of virion covered with the ch14D5 Fab fragments was generated by superimposition of the Fab-D3 complex structure onto D3 domains within cryo-EM structure of the mature TBEV virion (pdb_id 5O6V, Fuzik T et al., 2018). According to this model, Fab fragments can easily bind to type B and type C glycoprotein E molecules (Figure 3B). However, binding to type A molecules located close to the 5-fold rotation axes is hindered due to steric clashes between adjacent Fab molecules as well as between Fab molecules and glycoprotein E molecules (Figure S4). Thus, up to 120 molecules of the ch14D5 antibody Fab fragment can simultaneously bind to the mature TBEV virion at saturating Fab fragment concentrations (C >> K_D_). Moreover, intact ch14D5 antibody molecules are also able to bind to the virion as many as 120 copies per virion without overlapping (Figure 3C).

**Figure 3.**
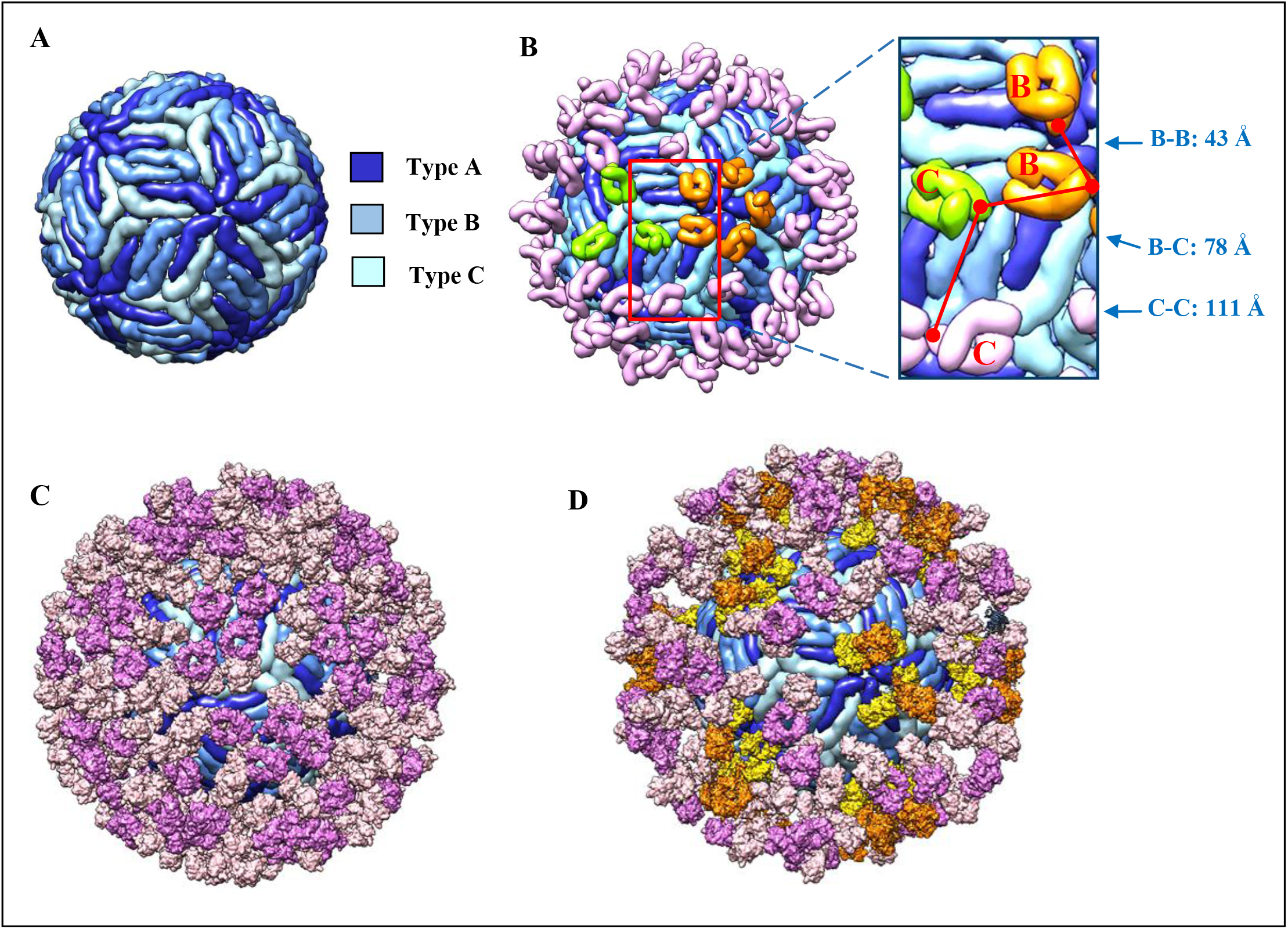
Models illustrating various possible complexes between the ch14D5 antibody (or its Fab fragment) and the mature TBEV virion. (A) Organization of glycoprotein E molecules on the surface of the mature TBEV virion. Glycoprotein E molecules close to 2-, 3-and 5-fold axes are shown in medium blue, light blue and deep blue, respectively. (B) Model of the Fab fragments of the ch14D5 antibody bound to the TBEV virion surface. Most of the Fab fragment molecules are shown in pink, while some of molecules bound to type B glycoprotein E molecules are shown in orange, and some of those bound to type C glycoprotein E molecules are shown in light green. (C and D) Models of the TBEV virion (shown in blue) coated by 120 molecules of the monovalently-bound ch14D5 antibody (C) or by both monovalently and bivalently-bound antibody molecules (D). Fab arms of the monovalently-bound antibodies are shown in light pink, while Fc fragments are shown in medium pink. Fab arms of bivalently-bound antibodies are shown in yellow, while Fc fragments are shown in orange.

We also checked whether the D1 and D2 domains of neighboring glycoprotein E molecules are also involved in antibody binding. We were unable to find direct evidence, since the distances between adjacent amino acid residues of the antibody and neighboring glycoprotein E were slightly higher than acceptable (Figure S5). However, given the high mobility of glycoprotein E molecules on the virion surface (Lok SM et al., 2008; Pierson TC et al., 2008) and the limited resolution of the virion model (about 3.9 Å according to Fuzik T et al., 2018), we believe that moderate interactions with neighboring glycoprotein E molecules are highly probable.

Further we checked whether the bivalent binding of the ch14D5 antibody to the virion is possible. We consider such binding to be theoretically possible if antibody molecule can simultaneously reach two adjacent epitopes on the surface of the virion. Based on known X-ray structures of full-size IgG antibodies, the maximum distance between heavy chain Lys218 C-alpha atoms was estimated to be approximately 45 Å, which is in good agreement with previous estimates (Edeling MA et al., 2014; Rippol DR et al., 2016). We then measured the distances between C-alpha atoms of Lys218 of adjacent Fab fragments in the model of the TBEV virion occupied by the ch14D5 Fab-fragments (Figure 3B). We took into account that antibodies are flexible (Zhang X et al., 2015; Jay JW et al., 2018), and that the angle between the VH/VL and CH1/CL domains (Fab elbow angle) can vary in a wide range (Stanfield RL et al., 2006; Kaufmann B et al., 2006). In the case of Fab fragments bound to type B glycoprotein E molecules (Figure 3B, orange molecules) the distance was 43 Å even for non-bended Fab fragments, whose conformation was taken directly from our X-ray structure (Figure 3B, inset). A slight rotation of the CH1/CL domains relative to the VH/VL domains around the longitudinal axis of the Fab fragment by 25° counterclockwise reduced this distance to 40 Å. In the case of the bivalent binding of two Fab arms of the same IgG1 molecule at binding sites of type B and C simultaneously, the distance between the corresponding Lys218 atoms would have to be 78 Å, which is far too big to be possible (Figure 3B). Bending one of the Fab fragments to a maximum possible elbow angle of 195° (Stanfield RL et al., 2006) led to a distance of 60 Å between the C-alpha atoms of Lys218 residues, which is also too big for the bivalent binding to occur. The bivalent binding of IgG1 molecule to two type C sites is impossible due to the same reason. Consequently, the bivalent binding of the ch14D5 antibody to type B glycoprotein E molecules is the only possible case. To further support this assumption, we generated a model of virion covered with 24 bivalently-bound ch14D5 molecules at type B positions (Figure 3D, orange-yellow IgG molecules) as well as 60 antibody molecules bound to type C glycoprotein E in a monovalent manner (Figure 3D, pink IgG molecules). The model satisfied the steric and geometric constraints without any clashes.

### *In silico* analysis of possible mechanisms of virus neutralization by the ch14D5 antibody at the post-attachment stage of cellular infection

The process of cell infection by TBEV comprises of two main stages, namely, binding to cell receptors, leading to endocytosis of the virion, and fusion of the viral membrane with the endosomal membrane during endosome maturation. Many successive steps accompany this process: pH-dependent dissociation of glycoprotein E dimers with the formation of monomers, rearrangement of the virion envelope, pH-dependent rotation of domains D1, D2 and D3 relative to each other, formation of pre-fusion glycoprotein E trimers and subsequent transition to fusion-competent trimers, displacement of the glycoprotein stem region, pulling of the endosomal membrane towards the viral one and the formation of a membrane pore (Figure 4A) (Ferlenghi I et al., 2001; Kuhsn RJ et al., 2002; Modis Y et al., 2004; Stiasny K et al., 2013; Chao LH et al., 2014; Zhang X et al., 2015, and also reviewed in Kaufmann B et al., 2011 and Pierson TC et al., 2013). We analyzed whether bound ch14D5 antibody or its Fab fragment could block some of these movements. First, we evaluated whether bivalently-bound antibody molecules can cross-link glycoprotein E molecules like a net and block viral envelope reorganization. According to the proposed mechanism (Kuhn RJ et al., 2002; Kaufmann B et al., 2006), the D3 domains of the type B glycoprotein E molecules have to move away from each other at a considerable distance (about 160 Å according to our estimates). At the same time, the distance between antigen-binding ends of the IgG molecule is limited to 180-190 Å (according to Zhang X et al., 2015, and also to our calculations for the 1HZH and 1IGT pdb structures). Given that the angle between the Fab arms of the ch14D5 antibody and the virion surface is about 40°, the bivalently-bound ch14D5 antibody will not allow the D3 domains to move away from each other by more than 140 Å (Figure 4B). Thus, if the proposed mechanism for the transition of the virion to the fusogenic state (Kuhn RJ et al., 2002) is correct, then the bivalently-bound ch14D5 antibody molecules should block or significantly slow down this process.

**Figure 4.**
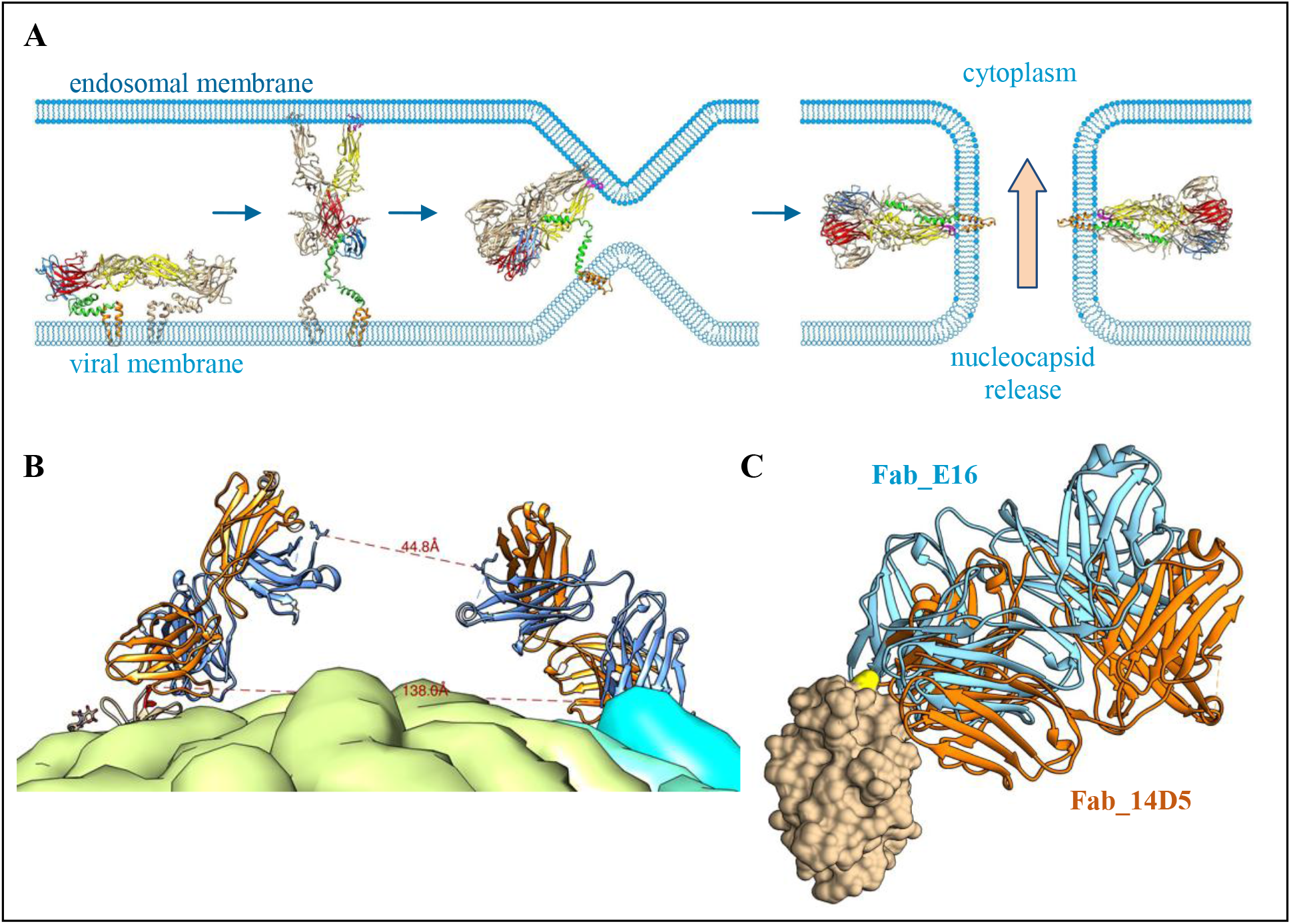
Evaluation of different scenarios of blocking the post-attachment steps of TBEV infection by the ch14D5 antibody. (A) The sequence of processes preceding release of the nucleocapsid into the cell cytoplasm. Transmembrane helices are shown in orange, fusion loop is shown in magenta. (B) A model of the TBEV virion (based on the 5A6O pdb structure) illustrating the maximum possible distance between D3 domains bound by the same antibody molecule. The right Fab fragment molecule, placed in the model artificially, was placed so that the distance between the C-alpha atoms of the Lys218 residues of the heavy chains for these two Fab fragments was about 45 Å, and the angle between the Fab fragment and the virus surface was equal to the binding angle for this antibody. (C) Two Fab fragments (14D5 is shown in orange while E16 is shown in light blue) superimposed onto the same D3 domain (shown in tan). The alignment was based on the 1ZTX pdb structure.

The ability to lock the virion requires that the antibody-antigen complex will be stable enough at acidic environment up to pH 5.0 (late endosomes). It was shown by SPR that the ch14D5 antibody binds to the D3_Sof protein with high affinity even at pH 4.8 (Figure S6).

It also has been previously shown that the E16 antibody and its Fab fragments block the infectivity of the West Nile virus at a post-attachment stage, presumably by interfering with the rearrangements of glycoprotein E domains (Kaufmann B et al., 2006; Kaufmann B et al., 2009). Comparison of the structures for Fab-D3 complexes of the ch14D5 or E16 antibodies showed that these Fab-fragments are oriented similarly (Figure 4C). Since the E16 antibody Fab fragment has been experimentally shown to block membrane fusion, it is likely that the ch14D5 Fab fragment, as well as the full-length ch14D5 antibody, will block membrane fusion in the same manner.

Thus, based on the orientation of the Fab fragment of the ch14D5 antibody relative to the D3 domain, and the current understanding of the geometric details of the intramolecular and intermolecular reorganization of glycoprotein E molecules, we propose that the full-length ch14D5 antibody can block not only viral binding to the cell, but also some steps of the intra-endosomal stage of infection.

## Discussion

In this study, the structure of the complex between the protective antibody ch14D5 and the D3 domain of TBEV glycoprotein E has been determined by X-ray diffraction. It has been shown that amino acid residues D308 – K315 and A331 – K336, as well as N367 and E387 residues located on the lateral ridge of the D3 domain of glycoprotein E, are involved in the interaction. These data explain the observed differences in affinity of the antibody to the D3 domains of Far-Eastern, Siberian and European virus subtypes. Structural analysis revealed that Ala331 residue of the D3 protein of Far-Eastern subtype is located in a small hydrophobic pocket formed by aromatic amino acid residues Tyr_L32 and Tyr_L50 (positions 32 and 50 of the antibody light chain), as well as Phe_H101 (heavy chain) (Figure 5). However, this pocket is too small for the amino acid residue Thr331 located in the same position in the Siberian and European D3 proteins. This presumably lead to destabilization of the complex and a decrease in affinity by 1-2 orders of magnitude.

**Figure 5.**
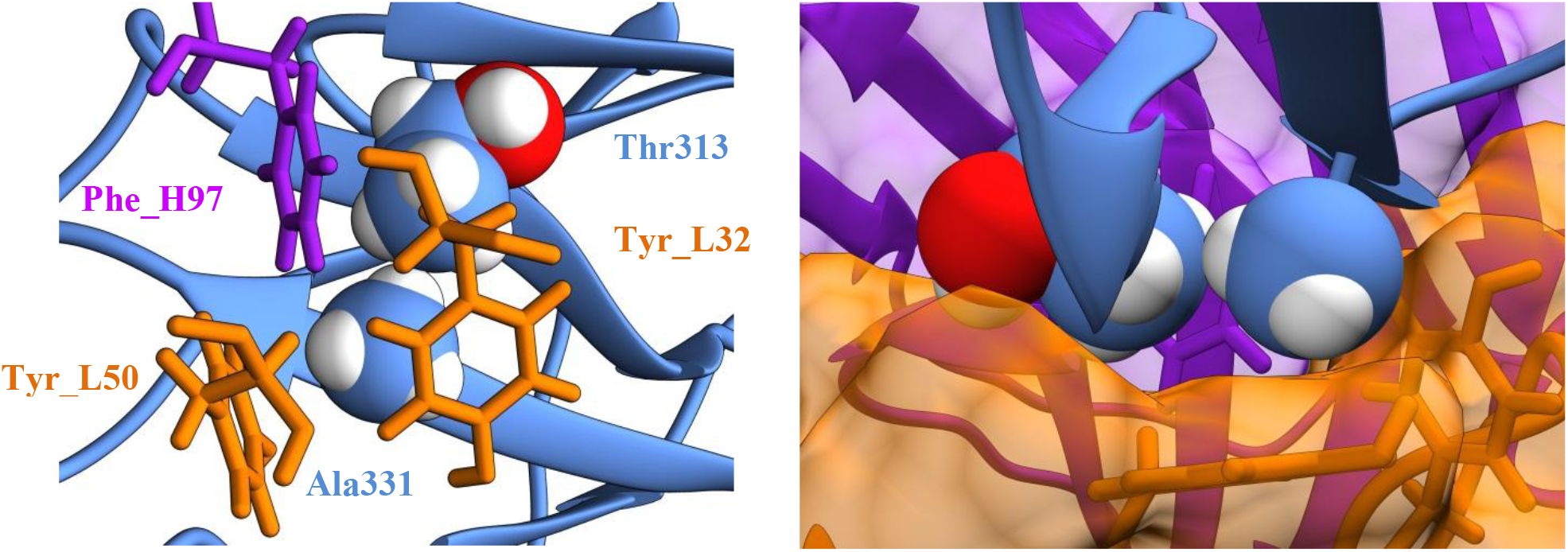
Structural reasons for the differences in binding of the ch14D5 antibody to recombinant D3 proteins. Structures illustrating tight neighborhood of Thr313 and Ala331 residues (shown as spheres). Heavy chain of antibody is shown in purple, light chain is shown in orange, the D3 domain is shown in blue.

The virus-neutralizing and protective activity of a particular antibody depends on its affinity as well as on the location of the epitope and its accessibility (reviewed in Dowd KA et al., 2011 and Austin SK et al., 2014). It has been shown that the epitope recognized by the ch14D5 antibody is readily accessible. Therefore, the binding of the ch14D5 antibody does not require additional time-consuming stages of “adjustment” of the virion surface, as in the case of antibodies directed to cryptic epitopes (Zhang X et al., 2015; Lok SM et al., 2008). Up to 120 molecules of the ch14D5 antibody or its Fab fragment can bind to the virion at saturating antibody concentrations, the number that is common for the majority of virus-neutralizing antibodies against flaviviruses (Kaufmann B et al., 2006; Fuzik T et al., 2018). It is believed that the higher this number, the higher the antiviral activity of neutralizing antibody (Pierson TC et al., 2007; Austin SK et al., 2014). Considering that even 60 accessible epitopes per virion are sufficient for successful neutralization (Fibriansah G et al., 2015), the presence of 120 accessible epitope copies, together with the high affinity of the ch14D5 antibody, is more than sufficient for successful neutralization of the virus and provides high protective properties of this antibody (Baykov IK et al., 2014). In addition, it was shown that the bivalent binding of the ch14D5 antibody to the virion is also possible. This usually results in a significant increase in the “functional” affinity (avidity) of antibody for the virus, as well as an increase in the neutralizing activity by 2−4 orders of magnitude (Wang P et al., 2010; Edeling MA et al., 2014; Plevka P et al., 2014; Ripoll DR et al., 2016).

Several mechanisms of antiviral activity for antibodies against flaviviruses are known (reviewed in Pierson TC et al., 2007; Dowd KA et al., 2011; Austin SK et al., 2014). Blocking the interaction of the virion with cellular receptors is one of them. TBEV can infect a variety of mammalian and tick cell types via various receptors and attachment factors. Laminin-binding protein and alphaVbeta3 integrin may be involved in the entry of TBEV virions into cells (Protopopova E et al., 1999; Zaitsev BN et al., 2014) as well as some receptors used by mosquito-borne flaviviruses, such as D-SIGN, TIM and TAM (reviewed in Laureti M et al., 2018; Yun SI et al., 2018). It is believed that the lateral ridge of the D3 domain of glycoprotein E is the main receptor binding site on the surface of the TBEV virion (reviewed in Mandl CW, 2005 and Kellman EM et al., 2018). This supported by the fact that the majority of highly neutralizing and protective mice-derived monoclonal antibodies and some human-derived antibodies against flaviviruses are directed to this region of the D3 domain (Roehrig JT, 2003; Oliphant T et al., 2005; Sánchez MD et al., 2005; Dai L et al., 2016; Agudelo M et al., 2021). Since the ch14D5 antibody recognizes the epitope located on the lateral ridge of the D3 domain, we assume that one of the major mechanisms of the antiviral action of this antibody is blocking the binding of the virion to cellular receptors.

It is likely that the ch14D5 antibody can also block some post-attachment steps of infection. Many details of this intra-endosomal stage of viral lifecycle are still unknown. However, it is known that this stage consists of several sequential steps, including the dissociation of glycoprotein E dimers to form monomers, the rearrangement of the virion envelope, the rotation of domains D1, D2 and D3 relative to each other, the formation of pre-fusion trimers, the transition of these structures to fusion-competent trimers, the formation of a membrane pore and, finally, the release of the nucleocapsid into the cell cytoplasm (Ferlenghi I et al., 2001; Kuhn RJ et al., 2002; Chao LH et al., 2014; Zhang X et al., 2015). Since the Fab arms of the ch14D5 antibody and highly potent antibody E16 against WNV have similar orientations relative to the glycoprotein E molecule, we believe that the ch14D5 antibody can block glycoprotein E rearrangement in a similar way as postulated for the E16 antibody (Kaufmann B et al., 2009). Since the ch14D5 antibody can also bind in a bivalent manner, this should block virion surface rearrangement by cross-linking glycoprotein E molecules. Thus, in addition to the blocking of virion attachment, two more mechanisms are highly possible for this antibody, namely, blocking through steric hindrance and through the cross-linking of glycoprotein E molecules.

We believe that blocking of infection at the post-attachment step may be particularly important for preventing antibody-dependent or antibody-mediated enhancement of infection (ADE or AME, respectively), which has previously been shown for various viruses (reviewed in Taylor A et al., 2015 and Tay MZ et al., 2019), including TBEV (Phillpotts RJ et al., 1985; Kreil TR et al., 1997) and other flaviviruses (Schlesinger JJ et al., 1981a and 1981b; Gollins SW et al., 1984; Ng JK et al., 2014; Ayala-Nunez NV et al., 2016; also reviewed in Dowd KA et al., 2011 and Halstead SB, 2014). While a virion opsonized by a sufficient number of neutralizing antibodies cannot infect regular cells, it can enter FcγR-bearing cells such as macrophages, monocytes and dendritic cells via FcγR-mediated phagocytosis. During the maturation of the phagosome, acidification of the internal space occurs, similar to acidification of the inner space of endosomes (reviewed in Kinchen JM et al., 2008 and Levin R et al., 2016). This acidification initiates conformational rearrangements in glycoprotein E molecules that lead to the formation of trimeric spikes with exposed fusion loops (Fritz R et al., 2008). If bound antibody molecules are unable to block membrane fusion, then the infection of FcγR-positive cells occurs, which leads to the formation of viral progeny. At some point, the number of protective/neutralizing antibodies introduced into the organism for pre-or post-exposure prophylaxis may become insufficient to reach the stoichiometric threshold required to successfully block all virions of the growing viral progeny (Pierson TC et al., 2007; Dowd KA et al., 2011; Austin SK et al., 2014). In this case, replication of the virus in regular cells can start, which can lead to systemic infection. However, if antibody molecules can catch or trap the virion in one of the intermediate states during the maturation of phagosomes thus blocking membrane fusion, then the phagosome has enough time to merge with lysosomes and form a phagolysosome. As a result, lysosomal enzymes destroy the virus (Levin R et al., 2016) and the infection does not develop. In the case of the ch14D5 antibody, it was shown that it is able to bind to the D3 domain at a reduced pH value (pH 4.8, which is even more acidic than pH ∼ 5.0 inside of late endosomes) almost as strongly as at physiological pH 7.2 (Figure S6). Our hypothesis that the ch14D5 antibody can block infection at the post-attachment step is in good agreement with the fact that this antibody, as well as the related chFVN145 antibody, did not induce ADE in *in vivo* experiments (Baykov IK et al., 2014; Matveev AL et al., 2019).

At the moment, experimental structures are only available for three antibodies against TBEV: 19/1786 (Fuzik T et al., 2018), Mab 4.2 (Yang X et al., 2019) and T025 (Agudelo M et al., 2021). Comparison of the structure determined in this study with these structures showed that the epitope of the ch14D5 antibody is adjacent and partially overlap with other epitopes, but it is not identical to them. Moreover, there are significant differences in the orientations of Fab fragments relative to the D3 domain and thus relative to the virion surface (Figures 6A and 6B). In particular, the angle between the longitudinal axes of the Fab fragments of the ch14D5 and 19/1786 antibodies is about 80°. The same is true for antibodies ch14D5 and T025, since antibodies 19/1786 and T025 have almost identical orientation (Agudelo M et al., 2021). The Fv fragment of antibody 4.2 takes an intermediate position, and it is rotated around the longitudinal axis by approximately 90°. In the context of the whole virion, this results in significantly different binding orientations (Figure 6B). Such differences can affect the possibility of bivalent binding for IgG molecules to the virion (Ripoll DR et al., 2016), which in turn affects the “functional” affinity and neutralizing properties of the antibody as well as molecular mechanism of action.

**Figure 6.**
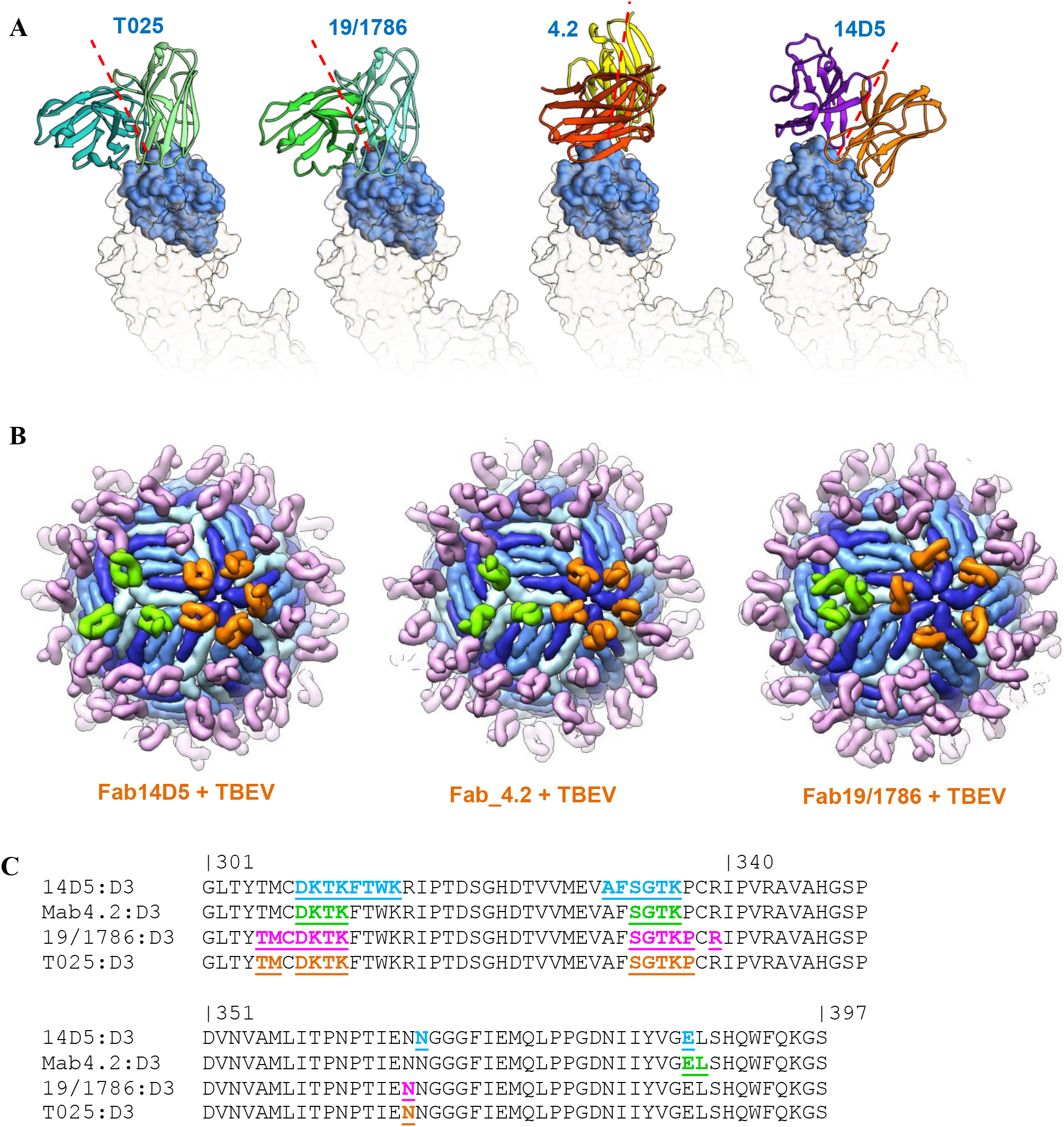
Differences between anti-TBEV antibodies 14D5, mab4.2, 19/1786 and T025 in terms of binding to glycoprotein E molecules. (A) Orientation of Fv fragments of antibodies 14D5, mab4.2, 19/1786 and T025 relative to the D3 domain of glycoprotein E. Orientation of the D3 domain is the same for all structures. A part of D1 and D2 domains is shown as transparent surface. Red dashed lines indicate longitudinal axes of corresponding Fab fragments. (B) Differences in orientation of Fab fragments of these antibodies in context of virion surface. Fab fragments bound to type B sites are shown in orange, while those bound to type C sites are shown in neon green. (C) Comparison of epitopes recognized by these antibodies. Residues are numbered according to Rey FA et al., 1995. Residues forming epitope are underlined and shown in color.

Since the epitopes of ch145, 19/1786, T025, and 4.2 antibodies overlap, these antibodies are likely to compete with each other when used simultaneously. Therefore, an increase in the protective activity with the simultaneous use of these antibodies is unlikely. At the same time, the combined use of these antibodies may be advisable, since such antibody “cocktail” may have a broader spectrum of action and may be more resistant to variations in the amino acid sequence of escape variants’ glycoprotein E. We have mapped the key amino acid residues of glycoprotein E that form the interface in each of the four complexes (Figure 6C). If a single substitution occurs in the amino acid sequence of these regions of glycoprotein E, it is possible that at least one of these antibodies will still effectively bind to the virus and block its spread.

We can conclude that the ch14D5 antibody recognizes a unique epitope on the lateral ridge of the D3 domain of glycoprotein E, and that the orientation of this antibody with respect to the viral glycoprotein E is unique among the antibodies against TBEV with a known structure. It was also shown that the epitope is spatially accessible, and that up to 120 IgG molecules or Fab fragments can bind to the virion simultaneously. Based on our findings, it can be concluded that there are two likely mechanisms of antiviral activity for this antibody, namely, blocking the interaction of the virion with cellular receptors and blocking membrane fusion inside endosomes or phagosomes. Our findings explain the high antiviral and protective activity of the ch14D5 antibody and the absence of ADE of infection. These data can be also used for structure-guided optimization of the properties of the antibody through rational design.

## Acknowledgements

Most of the experiments (including protein preparation, crystallization and structure determination), as well as the preparation of the manuscript and publication of the article, were carried out with the support of the Russian Science Foundation (grants 19-74-00107 and 17-74-10146). The preparation of recombinant proteins D3_Zau, D3_Vas, D3_Bal, and D3_Eu, as well as the analysis of the interaction of the ch14D5 antibody with these proteins, was supported by the Ministry of Science and Higher Education of the Russian Federation (grant of the President of the Russian Federation, project number MK-6575.2018.4). The authors would like to thank the organizers, teachers and tutors of the FEBS practical course “Advanced Methods in Macromolecular Crystallization VIII” (June 2018, Nove-Hrady, Czech Republic) for the opportunity to perform crystallization screening experiments and their valuable advice. We would also like to thank the researchers at EMBL Hamburg: Maria Marta Garcia Alai and Christian Günther for their assistance with the protein crystallization trials, Isabel Bento for valuable advice and Gleb Bourenkov for the assistance in using the PETRA III P14 beamline (EMBL, DESY, Hamburg). We would like to thank the researchers at ICBFM SB RAS, Novosibirsk: Artem Yu. Tikunov and Igor V. Babkin for their assistance in the DNA sequencing, along with Lyudmila M. Sokolova, Galina B. Kaverina, Vladimir F. Podgorny, and Alexei A. Ilyichev for their assistance in the production of D3 proteins in preparative quantities, as well as the students Evgeniya Karelina, Pavel Desyukevich, Ekaterina Mikhaylova, and Olga Kurchenko for their help with some experiments.

## Author contributions

I.K.B. and N.V.T. conceived and designed the study. I.K.B. and L.A.E. contributed to protein preparation and purification. A.L.M. produced the ch14D5 antibody. I.K.B. performed protein crystallization with assistance from N.A.M. G.C., I.K.B., P.P. and P.M.R. contributed to X-ray data collection, structure solution and refinement. I.K.B. preformed binding experiments and 3D modelling. V.S.L. provided support and supervision of experiments at the EMBL, Hamburg, Germany. N.V.T. provided support and supervision of experiments at the Institute of Chemical Biology and Fundamental Medicine, SB RAS, Novosibirsk, Russia. I.K.B. and N.V.T. combined the data and wrote the manuscript. All authors reviewed and approved the pre-print version of the manuscript.

## Materials and methods

### Protein production and purification

Recombinant protein D3_Sof representing the D3 domain of the TBEV glycoprotein E, strain Sofjin-Ru, was obtained by periplasmic bacterial expression using pHEN2-rED3_301 plasmid (Baykov IK et al., 2018) and *E. coli* strain HB2151. The bacteria were cultivated in 3 liters of LB medium supplemented with 100 μg/ml ampicillin and 0,1% glucose in a Sartorius “BIOSTAT A plus” fermenter equipped with 5L vessel at 37 ° C, 200 rpm agitation and 3 L/min air flow rate. When the optical density OD_600_ reached 0.7–0.9, protein synthesis was induced by adding isopropyl-β-D-1-thiogalactopyranoside (IPTG) to a final concentration of 1 mM; the cultivation was continued at 30 °C for 4 h. Periplasmic proteins extraction and metal chelate chromatography were performed as described previously (Baykov IK et al., 2018). The protein was additionally purified by ion-exchange chromatography on a “POROS HQ/20” 4.6×100 mm column (Applied Biosystems) using 0-20%B gradient over 12 column volumes at 0.5 ml/min flow rate using 20 mM Tris-HCl buffer (pH 8.9 at 25 °C) as buffer A and the same buffer supplemented with 0.5M NaCl as buffer B. The protein was buffer-exchanged into phosphate-buffered saline supplemented with 0.02% sodium azide (PBSN) pH 7.2 and concentrated to a 2-4 mg/ml concentration using Amicon Ultra-4, 3 kDa cut-off (Millipore).

The Fab fragment of the ch14D5 antibody was prepared from the antibody using Immobilized Papain-agarose resin (Thermo Fisher Scientific) according to the manufacturer’s recommendations. It was then separated using Protein-A agarose resin (Thermo Fisher Scientific). The protein was additionally purified by ion-exchange chromatography on a “Mono S 5/50 GL” column (GE healthcare) at 0-100% B gradient over 15 column volumes at 0.5 ml/min flow rate using 20 mM sodium phosphate buffer (pH 6.0 at 25 °C) as buffer A and 20 mM sodium phosphate buffer (pH 7.0 at 25 °C) as buffer B. The protein was buffer-exchanged into PBSN buffer and concentrated to a 2-4 mg/ml concentration using Amicon Ultra-4, 30 kDa cut-off (Millipore). The purity of the obtained proteins was confirmed by sodium dodecyl sulphate–polyacrylamide gel electrophoresis.

### Protein crystallization

The protein complex was formed by incubation of the purified D3_Sof protein and the Fab-fragment of the ch14D5 antibody in a mass ratio of 3:8 (1.5x molar excess of D3_Sof protein) in a buffer containing 150 mM NaCl and 25 mM Tris-HCl (pH 7.5 at 25 °C) at room temperature for 1 h. The complex was separated from free D3_Sof protein by size-exclusion chromatography using “Superdex 75 10/300 GL” column (GE Healthcare) at 0.4 ml/min flow rate using running buffer containing 25 mM Tris-HCl (pH 7.6 at 25 ° C), 150 mM NaCl and 0.02% sodium azide. It was then concentrated to a 15 mg/ml concentration using Amicon Ultra-4, 3 kDa cut-off (Millipore).

The protein complex was crystallized by the hanging drop vapor diffusion method in 24-well plates (Hampton Research) at 25 ° C using an optimized precipitant. The precipitation solution was prepared by mixing 50 μl of EDO_P8K solution (Molecular Dimensions), 4.5 μl of 1 M imidazole solution (Molecular Dimensions), 5.5 μl of 1 M MES solution (Molecular Dimensions), 50 μl of solution number 33 from PEG/Ion screen (Hampton Research) and 40 μl H_2_O. 3 μl of the protein complex solution was mixed with 2 μl of the precipitant solution. 1.5 M NaCl solution was used as the reservoir solution. Hexagonal rod-shaped crystals ∼ 200 μm long with sharp tips formed after 7 days. Single crystals were harvested and cryo-cooled in liquid nitrogen for data collection.

### X-ray diffraction data collection

X-ray data were collected at the PETRA III storage ring (DESY, Hamburg, Germany), at EMBL beamline P14 using EIGERX 16M detector (Dectris, Baden, Switzerland). The data were indexed and integrated using XDS (Kabsch W, 2010) and scaled with AIMLESS (Evans PR et al., 2013) to 2.3 Å resolution. The crystal structure was solved by molecular replacement using MoRDa (Vagin A et al., 2015). The model was manually using COOT (Emsley P et al., 2010) and refined using REFMAC (Murshudov GN et al., 2011) in CCP4Cloud v1.6.020 (Krissinel E et al., 2018). The X-ray data collection and structure refinement statistics is presented in Table S1. The quality of the structures was assessed with the MolProbity server and wwPDB validation service.

### Interface analysis

The structure of the complex and interacting amino acids were analyzed using UCSF Chimera version 1.13.1 and PDBe PISA v1.52 (Krissinel E and Henrick K, 2007).

### Generation of D3 protein variants

The D3 proteins representing the D3 domains of the TBEV glycoprotein E belonging to various subtypes were used in this study. Plasmid DNAs pHEN2-rED3_301 (Baykov IK et al., 2018), pHEN2-D3_ZauM, pHEN2-D3_Eu and pHEN2-D3_Bal (Baykov IK et al., 2019) were used to produce proteins D3_Sof (Sofjin-Ru strain of TBEV), D3_ZauT331M (TBEV-2781 strain), D3_Eu (Absettarov strain of European subtype) and D3_Bal (1528-99 strain of Baltic lineage of Siberian subtype), respectively. In order to produce D3 proteins of Vasilchenko-like TBEV strain, one strain of Bosnia lineage of TBEV and one strain of Omsk hemorrhagic fever virus (OHFV), corresponding plasmid DNAs were constructed based on cDNA fragments of these strains: TBEV-356ICR (GenBank: MG598849.1), Bosnia-3 (GenBank: MH645616.1) and strain 186 of OHFV (Tkachev S. E., personal communication). The corresponding PCR-produced DNA fragments were inserted into the pHEN2 plasmid using *Nco*I/*Not*I endonucleases as described previously (Baykov IK et al., 2019). Overlap extension PCR mutagenesis was performed to obtain plasmid pHEN2-D3_SofA331T using primers D3_A331T_dir: 5’-GTCATGGAAGTTACCTTCTCTGGGACC-3’ and D3_A331T_rev: 5’-GGTCCCAGAGAAGGTAACTTCCATGAC-3’ and the plasmid DNA pHEN2-rED3_301 as a matrix for PCR.

The D3 proteins were produced similarly to the D3_Sof protein. However, smaller culture volumes of 600 ml were used for each protein. Periplasmic proteins extraction and metal chelate chromatography were performed similarly to described for the D3_Sof protein, ion-exchange chromatography was not used. The proteins were buffer-exchanged into PBSN buffer concentrated to 2-4 mg/ml concentration using Amicon Ultra-4, 30 kDa cut-off (Millipore). The purity of the obtained proteins was confirmed by sodium dodecyl sulphate–polyacrylamide gel electrophoresis.

### Affinity constant measurements

The binding of the ch14D5 antibody to the D3 domain variants was determined by a surface plasmon resonance-based biosensor assay (SPR), using a ProteOn XPR36 system (Bio-Rad Laboratories, Hercules, CA USA). PBS supplemented with 0.005% Tween-20 and 0.1 mM ethylenediaminetetraacetic acid (EDTA) was used as a system running buffer. The temperature of the sensor chip and autosampler was kept at 25 °C. Two vertical channels of HTG sensor chip were activated with 1 mM NiCl_2_ solution for 120 s at 30 μl/min flow rate. One of the D3 proteins was used for immobilization onto one of the activated channels resulting in a 50−80 response units (RU) immobilization level. Serial three-fold dilutions of the ch14D5 antibody in system running buffer starting from 81 nM (in the case of D3_Sof, D3_ZauT331M and D3_VasFSVA proteins) or 405 nM (in the case of D3_Eu, D3_Bal, D3_Bos, and D3_SofA331T proteins) concentrations were sequentially analyzed at a 25 μL/min flow rate at horizontal orientation of the micro channel module (MCM). These concentration ranges were determined based on data from scouting experiments. The same spot of sensor chip was used sequentially for all the measurements within the same group in order to exclude any immobilization level inconsistency. The duration of both the association and dissociation phases was 300 s. The chip surface was regenerated between each measurement by passing 300 mM EDTA solution for 120 s, followed by surface re-activation and D3 protein re-immobilization. The signal measured at the part of the chip surface where no D3 protein was immobilized was used as a reference. Global analysis of experimental data based on a single-site model was performed using the ProteOn Manager v. 3.1.0 software. The measurements were repeated at least three times, and average dissociation constants (K_D_) and their standard deviations (SD) were determined.

For pH-dependency measurements, D3_Sof protein was immobilized covalently on the surface of one of the vertical channels of the GLC sensor chip (Bio-Rad Laboratories, Hercules, CA USA) to a 100 RU level. Immobilization procedure was performed according to the manufacturer’s recommendations: carboxyl groups of alginate matrix were activated using a mixture containing equal amounts of freshly prepared 10 mM N-hydroxysulfosuccinimide and 40 mM 1-Ethyl-3-(3-dimethylaminopropyl)carbodiimide. A solution of D3_Sof protein at 5.5 μg/mL concentration in 10 mM sodium acetate buffer, pH 4.5, was used for immobilization. As soon as desired immobilization level was achieved, unreacted activated carboxyl groups of chip surface were quenched with 1 M ethanolamine-HCl, pH 7.5. Either PBS supplemented with 0.005% Tween-20 or 100 mM sodium citrate buffer (pH 4.8 at 25 °C) supplemented with 150 mM NaCl and 0.005% Tween-20 was used as a system running buffer. Serial three-fold dilutions of the ch14D5 antibody in system buffer starting from 81 nM were sequentially analyzed at a flow rate of 25 μL/min and horizontal orientation of the MCM.

The duration of both the association and dissociation phases was 300 s. The chip surface was regenerated by passing 50 mM NaOH solution for 40 s between each measurement. The signal measured at the part of the chip surface where no D3 protein was immobilized was used as a reference. Global analysis of experimental data based on a single-site model was performed using the ProteOn Manager v. 3.1.0 software. The measurements were repeated three times, and average dissociation constants and their standard deviations were determined.

### Modelling and structural analysis of antibody-covered TBEV virion

The cryo-EM derived TBEV virion model (the 5O6A pdb structure) was used to build a virion model coated with a the ch14D5 antibody Fab fragments or the whole ch14D5 antibody molecules. MatchMaker tool from UCSF Chimera v. 1.13.1 was used to align the D3 domain of the Fab-D3 complex with the D3 domain of glycoprotein E molecules (chains B and C of the 5O6A model). “Multiscale models” tool from Chimera with icosahedral symmetry selected was used to generate virion model from asymmetric unit. In order to create a model of a virion occupied by the monovalently-bound full-length ch14D5 antibody, variable domains of IgG antibody (the 1HZH pdb structure) were fitted onto corresponding domains of the ch14D5 Fab fragment within a model of virion covered with Fab fragments. In order to build a model of virion with bivalently-bound antibody molecules, manual bending of peptide chain within hinge region of antibody was used.

### Comparison of structures of antigen-antibody complexes

MatchMaker tool from UCSF Chimera v. 1.13.1 was used for structural alignment of the D3 domains of various Fab-D3 complexes.

## Supplemental Information

**Table S1.** X-ray data collection and structure refinement statistics. The values for outer resolution shell are given in parentheses.

| **Data collection** |  |
| --- | --- |
| Space group | P 6<sub>1</sub> |
| a, b, c [Å] | 115.37, 115.37, 76.73 |
| $\alpha$, $\beta$, $\gamma$ [degrees] | 90, 90, 120 |
| Resolution [Å] | 60.9 – 2.30 (2.38 – 2.30) |
| Completeness [%] | 95.5 (99.9) |
| Wavelength [Å] |  |
| Multiplicity | 10.1 (10.2) |
| I/$\sigma$(I) | 17.9 (2.0) |
| CC1/2 [%] | 99.9 (80.4) |
| Rmerge [%] | 6.6 (119.0) |
| **Refinement statistics** |  |
| Rwork/Rfree [%] | 20.4/24.8 |
| Wilson plot B-factor [Å<sup>2</sup>] | 59.5 |
| R.m.s. deviations from target stereochemistry |  |
| Bond lengths [Å] | 0.01 |
| Bond angles [degrees] | 1.99 |
| Ramachandran plot [%] |  |
| Favored | 97.0 |
| Allowed | 2.8 |
| Outliers | 0.2 |
| MolProbity clashscore | 5 |

**Table S2.**
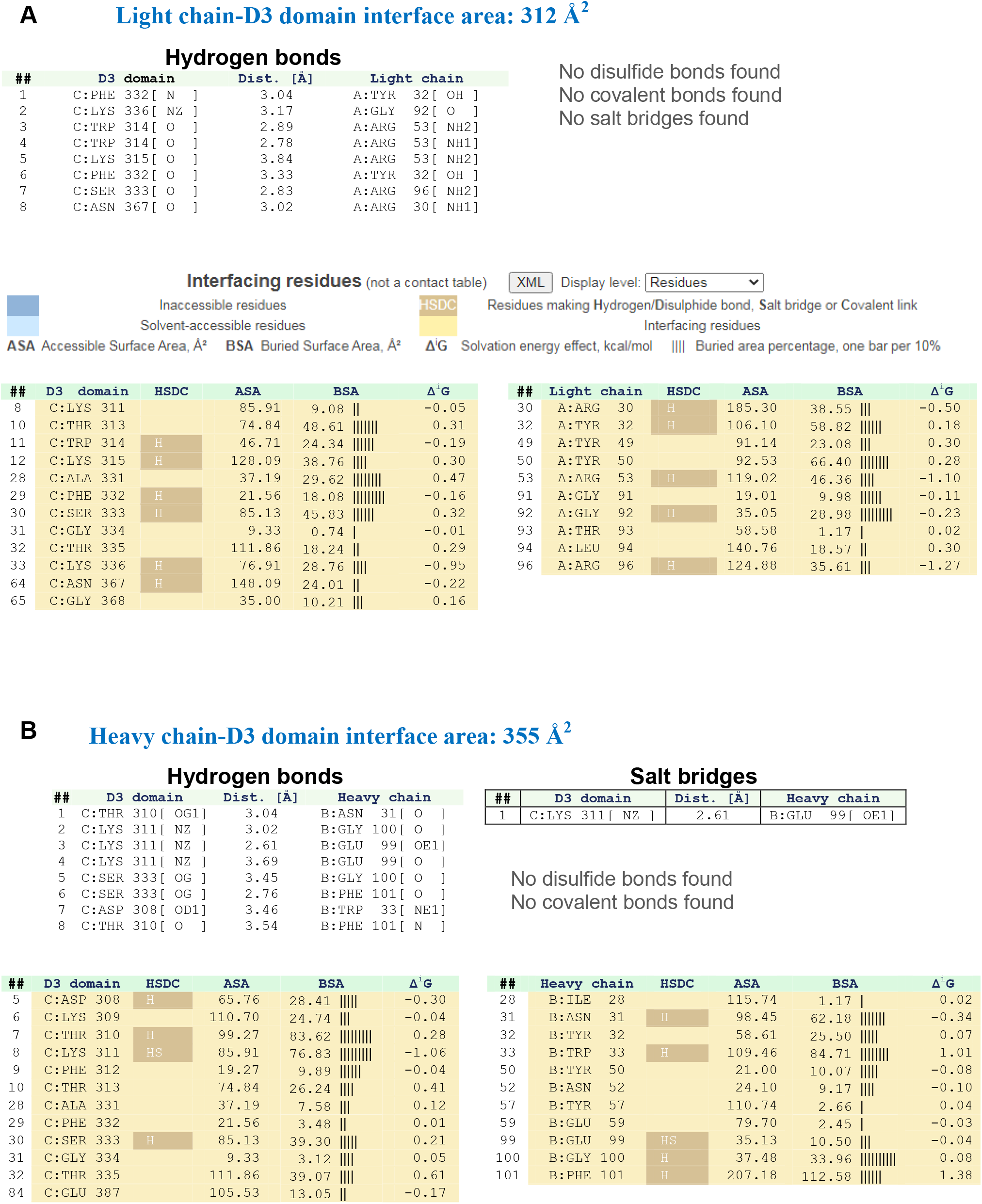
Table from PDBePISA interface analysis.

**Figure S1.**
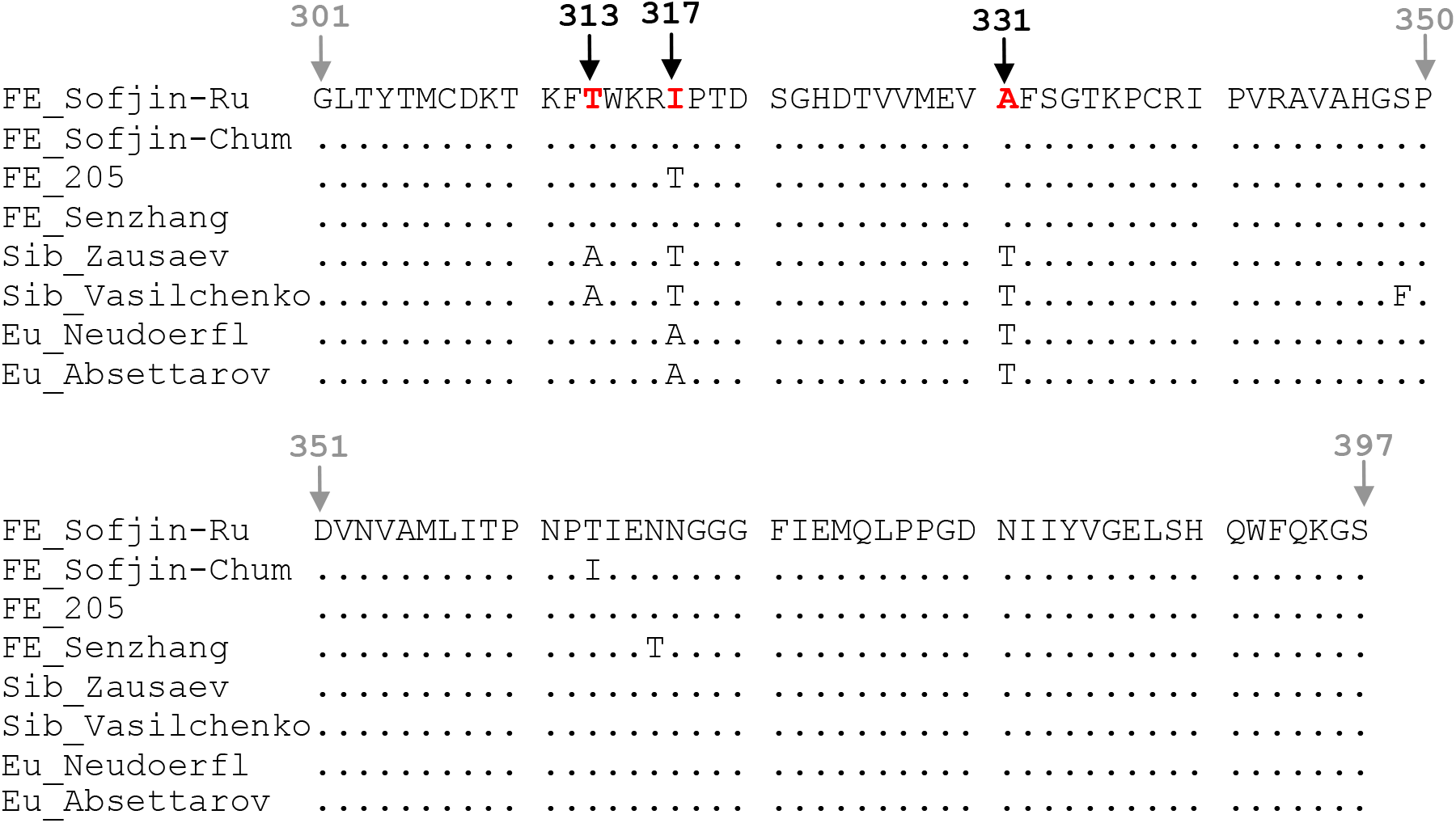
Alignment of amino acid sequences of glycoprotein E D3 domains belonging to the Far-Eastern, Siberian and European subtypes of TBEV. Positions for subtype-specific amino acid residues are in red.

**Figure S2.**
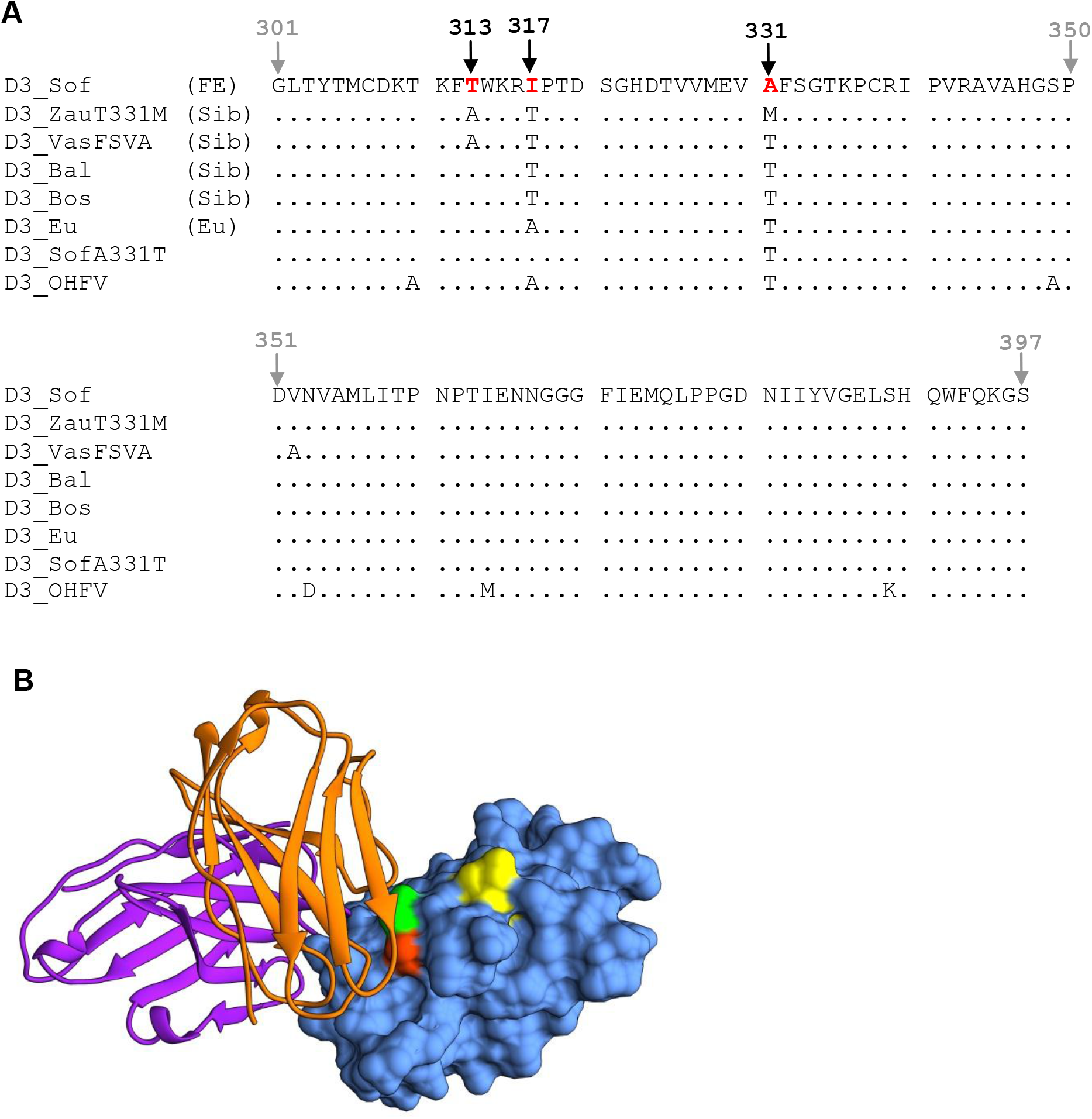
Amino acid variations between different strains of TBEV. (A) Alignment of amino acid sequences of glycoprotein E D3 domains used for affinity measurements. Positions for subtype-specific amino acid residues are in red. (B) Structure of the complex formed by the ch14D5 antibody Fv fragment and the D3 domain with residues 313, 317 and 331 colored in neon green, yellow and orange-red, respectively.

**Figure S3.**
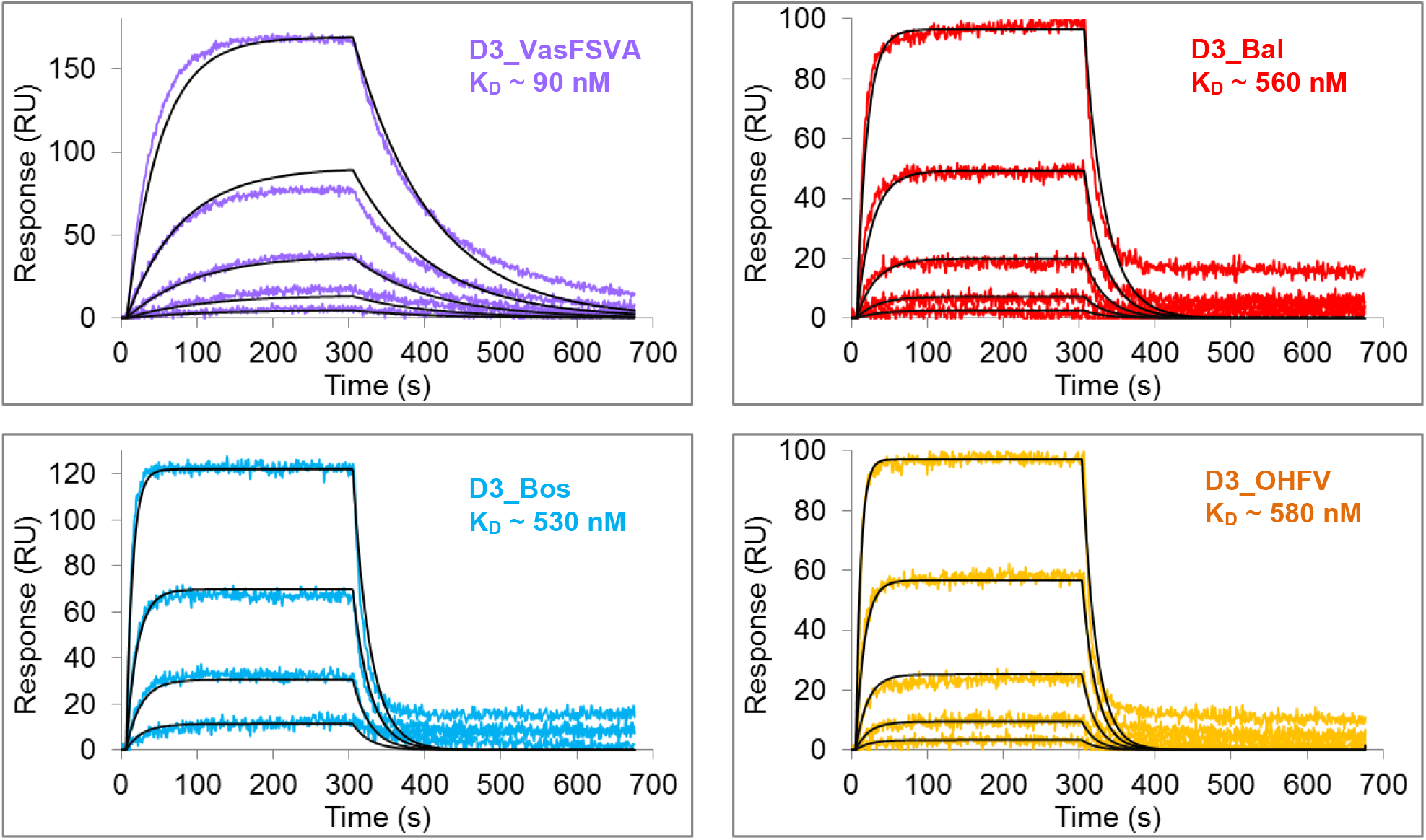
Differences in binding of the ch14D5 antibody to recombinant D3 proteins D3_Bal, D3_Bos, D3_ZauT331M and D3_VasFSVA, D3_OHFV detected by SPR. The D3 proteins were immobilized onto biosensor chip surface, the ch14D5 antibody was used as three-fold dilutions starting from 81 nM concentration (in the case of D3_VasFSVA protein) or from 405 nM (in the case of D3_Bal, D3_Bos and D3_OHFV proteins). Experimental curves are shown by colored lines, approximations are shown in black. The analysis was carried out using a ProteOn XPR36 biosensor.

**Table S3.** ProteOn XPR36 kinetic and affinity data.

| D3 protein | $k_{\text{on}}$, mean ± SD, $\text{M}^{-1}\text{s}^{-1}$ | $k_{\text{off}}$, mean ± SD, $\text{s}^{-1}$ | Average equilibrium dissociation constant $K_{\text{D}}$, mean ± SD, nM |
| --- | --- | --- | --- |
| D3_Sof | $1.1 \pm 0.2 \times 10^5$ | $3.3 \pm 1.7 \times 10^{-4}$ | $3.0 \pm 1.4$ |
| D3_ZauT331M | $8.0 \pm 2.4 \times 10^5$ | $1.1 \pm 0.2 \times 10^{-2}$ | $15 \pm 7$ |
| D3_VasFSVA | $1.4 \pm 0.2 \times 10^5$ | $1.2 \pm 0.2 \times 10^{-2}$ | $88 \pm 27$ |
| D3_Eu | $9.9 \pm 0.2 \times 10^4$ | $3.2 \pm 0.9 \times 10^{-2}$ | $320 \pm 90$ |
| D3_Bal | $9.7 \pm 1.2 \times 10^4$ | $5.5 \pm 2.1 \times 10^{-2}$ | $560 \pm 190$ |
| D3_Bos | $1.9 \pm 1.1 \times 10^5$ | $8.1 \pm 0.3 \times 10^{-2}$ | $530 \pm 300$ |
| D3_SofA331T | $9.7 \pm 0.2 \times 10^4$ | $3.1 \pm 0.8 \times 10^{-2}$ | $330 \pm 80$ |
| D3_OHFV | $1.6 \pm 1.0 \times 10^5$ | $6.8 \pm 0.9 \times 10^{-2}$ | $580 \pm 320$ |

**Figure S4.**
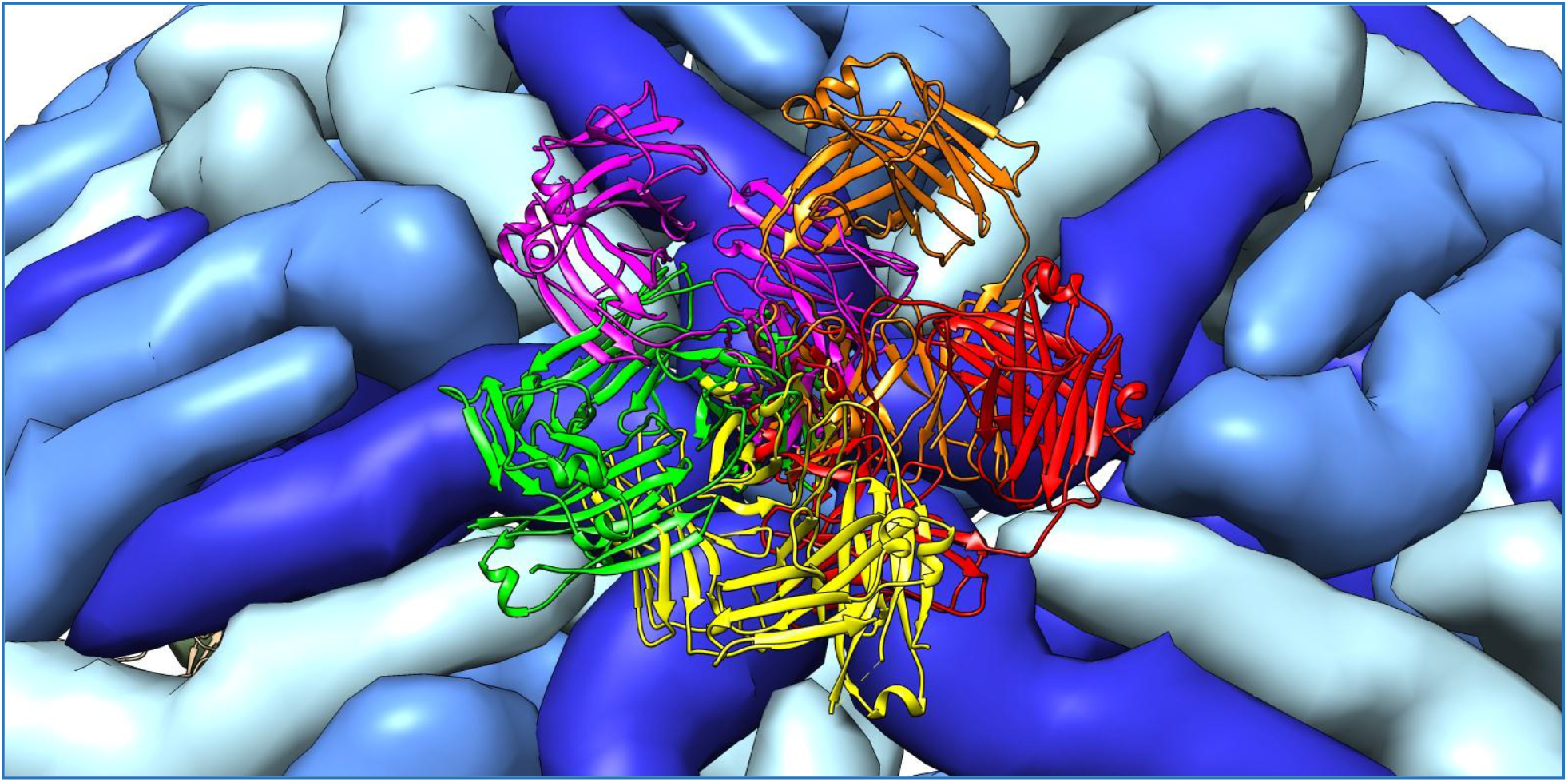
Clashes between the ch14D5 antibody Fab fragments bound to type A glycoprotein E molecules (near 5-fold rotation axis) Fab fragment molecules are shown in green, purple, orange, red and yellow. Type A glycoprotein E molecules are shown in deep blue.

**Figure S5.**
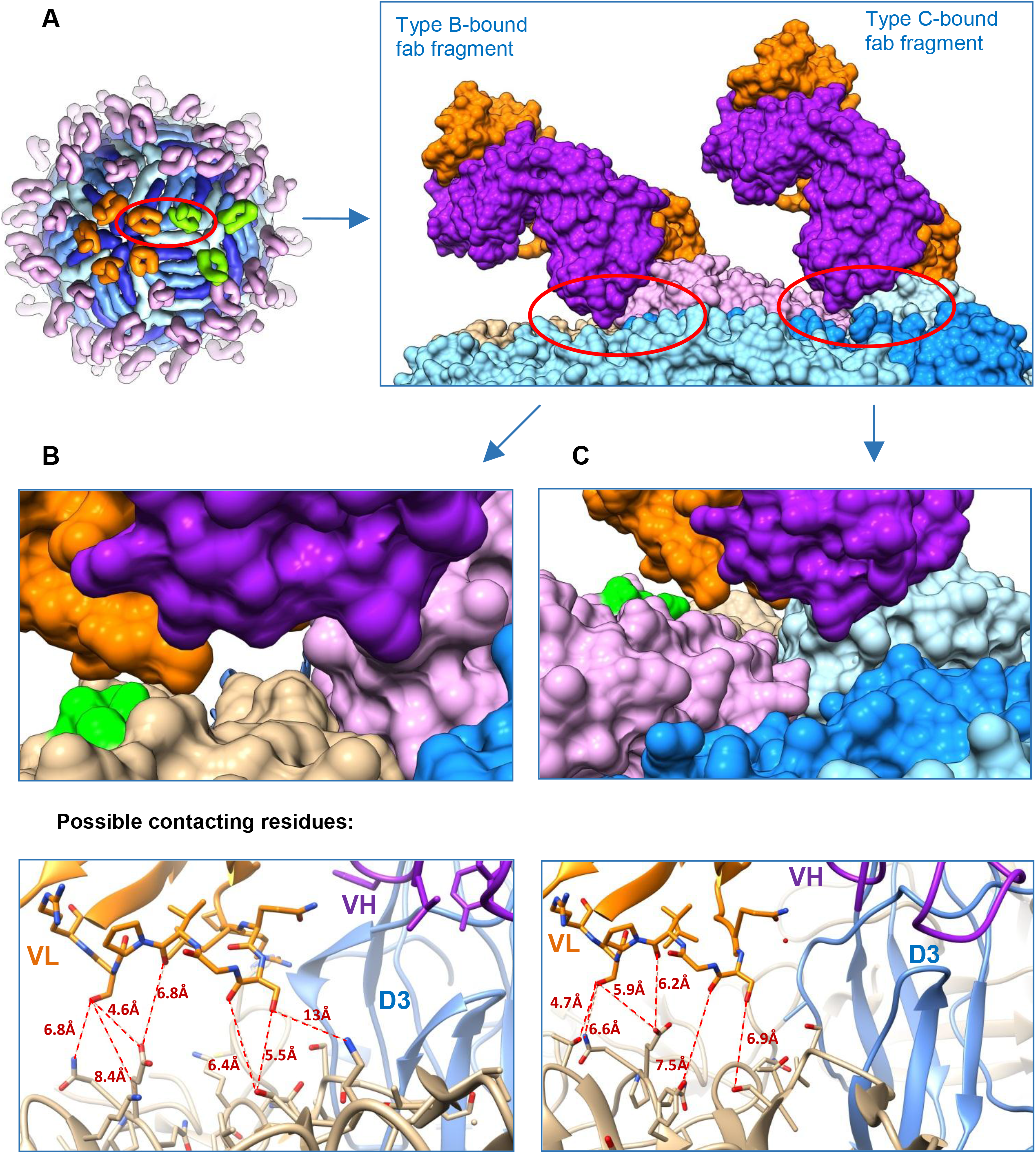
Involvement of adjacent glycoprotein E molecules into binding of the ch14D5 antibody to the virion. (A) Overall view (B) Close-up view of contacts between Fab molecule (bound to type B glycoprotein E molecule, shown in pink) and neighboring type A molecule of glycoprotein E, shown in tan (contacting surface is shown in light green). (C) Close-up view of contacts between Fab molecule (bound to type C glycoprotein E molecule, shown in light blue) and neighboring type B molecule of glycoprotein E, shown in pink (contacting surface is shown in light green).

**Figure S6.**
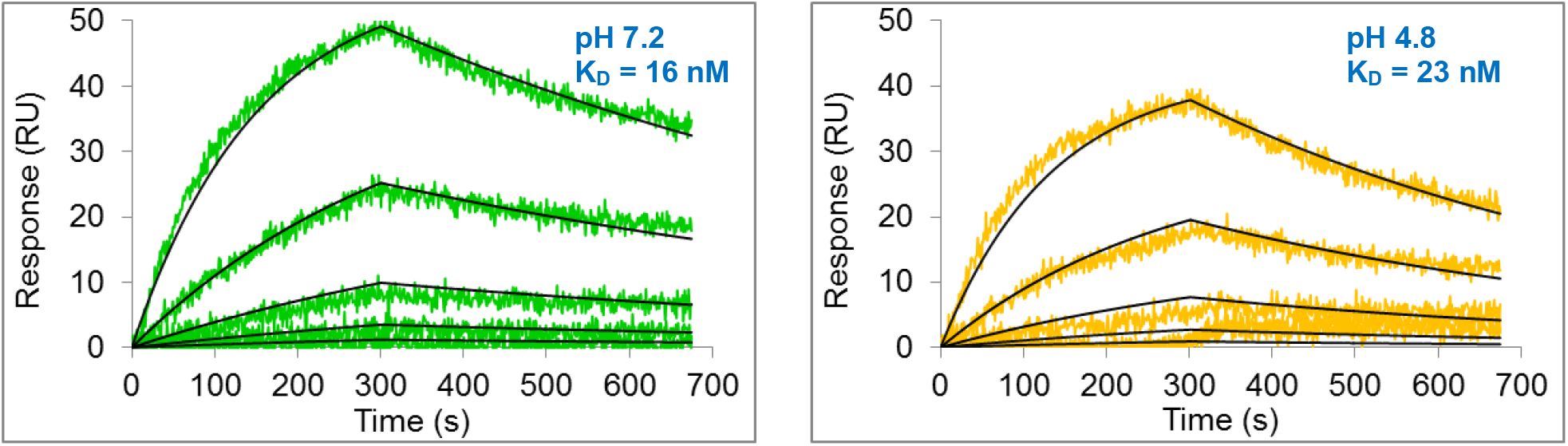
Binding of the ch14D5 antibody to the D3_Sof protein at pH 7.2 vs. pH 4.8. D3_Sof protein was immobilized onto GLC chip surface, serial three-fold dilutions of the ch14D5 antibody in running buffer starting from 81 nM concentration were analyzed. Reference-subtracted experimental data are shown as color lines, fitted data are shown in black.

